# High fiber, whole foods dietary intervention alters the human gut microbiome but not fecal short-chain fatty acids

**DOI:** 10.1101/2021.02.04.429869

**Authors:** Andrew Oliver, Alexander B. Chase, Claudia Weihe, Stephanie B. Orchanian, Stefan F. Riedel, Clark Hendrickson, Mi Lay, Julia Massimelli Sewall, Jennifer B. H. Martiny, Katrine Whiteson

## Abstract

Dietary shifts can have a direct impact on the gut microbiome by preferentially selecting for microbes capable of utilizing the various dietary nutrients. Intake of dietary fiber has decreased precipitously in the last century, while consumption of processed foods has increased. Fiber, or microbiota-accessible carbohydrates (MACs), persist in the digestive tract and can be metabolized by specific bacteria encoding fiber degrading enzymes. Digestion of MACs results in the accumulation of short-chain fatty acids (SCFAs) and other metabolic byproducts that are critical to human health. Here, we implemented a two-week dietary fiber intervention aiming for 40-50 grams of fiber per day within the context of a course-based undergraduate research experience (CURE) (n = 20). By coupling shotgun metagenomic sequencing and targeted gas-chromatography mass spectrometry (GC/MS), we found that the dietary intervention significantly altered the composition of individual gut microbiomes, accounting for 8.3% of the longitudinal variability within subjects. Notably, microbial taxa that increased in relative abundance as a result of the diet change included known MAC degraders (i.e., *Bifidobacterium* and *Lactobacillus*). We further assessed the genetic diversity within *Bifidobacterium*, assayed by amplification of the *groEL* gene. Concomitant with microbial composition changes, we show an increase in the abundance of genes involved in inositol degradation. Despite these changes in gut microbiome composition, we did not detect a consistent shift in SCFA abundance. Collectively, our results demonstrate that on a short-term timescale of two weeks, increased fiber intake can induce compositional changes of the gut microbiome, including an increase in MAC degrading bacteria.

**IMPORTANCE:** A profound decrease in the consumption of dietary fiber in many parts of the world in the last century may be associated with the increasing prevalence of Type II diabetes, colon cancer, and other health problems. A typical U.S. diet includes about ∼15 grams of fiber per day, far less fiber than daily recommended allowance. Changes in dietary fiber intake affect human health not only through the uptake of nutrients directly, but also indirectly through changes in the microbial community and their associated metabolism. Here we conducted a two-week diet intervention in healthy young adults to investigate the impact of fiber consumption on the gut microbiome. Participants increased their average fiber consumption by 25 grams/day on average for two weeks. The high fiber diet intervention altered the gut microbiome of the study participants, including increases in known fiber degrading microbes such as *Bifidobacterium* and *Lactobacillus*.

## INTRODUCTION

Consumption of dietary fiber has declined dramatically in the last century as processed foods have become a larger part of diets in the industrialized world. Pre-industrial and modern-day rural societies consume between 60-120 grams (g)/day of fiber, while individuals in the United States consume about half of the daily recommended allowance of 38 g/day for men and 25 g/day for women (1, 2). Declines in fiber intake over the past century have contributed to complications for human health. For example, chronic low fiber intake has been associated with Type 2 diabetes mellitus, heart disease, and colon cancer (3–5). Indeed, a reciprocal diet intervention exchanging African Americans low-fiber western diet with rural Africans high-fiber diet (increasing on average 40g per day) led to significant decreases in pre-cancerous biomarkers, further providing a link between fiber and human health (6). Furthermore, dietary fiber has been shown to protect against influenza infection (7), and may influence vaccine efficacy (8).

Dietary fiber is a mixture of polysaccharides that resist rapid digestion in the small intestine by endogenous enzymes and persists through the digestive tract into the colon. Once in the colon, fiber can be digested by the resident microbes (1, 9). This is due, in part, to the human genome encoding only 17 enzymes (i.e., glycoside hydrolases) that are capable of digesting carbohydrates (10). Conversely, the resident gut microbial communities collectively encode thousands of diverse enzymes from 152 gene families that can break down dietary fiber (11). In the colon, specialized microbes metabolize recalcitrant carbohydrates and produce fermented byproducts, including short chain fatty acids (SCFAs) such as acetate, propionate, and butyrate (12). SCFAs are capable of being absorbed across the human intestinal epithelial cells, and have direct impacts on human health (reviewed in (13)) such as stimulating and maintaining the mucus layer for the gut epithelium (14) and providing an energy source for butyrate-consuming colonocytes (15). SCFAs have also been shown to have immunomodulatory effects, including increased viral protection through altered T-cell metabolism (7), and inhibitory effects on pathogenic bacteria (e.g. *Clostridioides difficile*) (16).

Understanding the role of dietary fiber in structuring the gut microbiota could provide insights into managing chronic diseases associated with the gut microbiome. Typical diet intervention studies assessing the impact of fiber on gut microbial communities and the production of SCFAs have relied on single fiber supplements (17–19). Fiber supplements such as psyllium husks, inulin, wheat bran, resistant potato starch, and resistant corn starch vary in their efficacy for each individual (17, 20). Individuals might be more or less susceptible to the intervention depending on their initial resident microbial community and its ability to digest a particular fiber supplement. For example, one group investigating the impact of three fermentable fibers on gut microbiome composition and SCFA abundance found no significant effect when study participants consumed 20-24g resistant maize starch per day for two weeks (17). However, in addition to the quantity, the variety of dietary fibers may be important. Studies that have increased dietary fiber have previously observed changes in microbiome composition (3, 17, 18), yet results remain mixed on SCFA production (6, 21, 22). Further, the American Gut Project found that individuals who eat more than 30 types of plants in a week have a more diverse gut microbiome (23). Thus, the consumption of a diversity of fiber sources through whole foods may provide more opportunities for an individual’s gut microbiome to respond to the dietary changes and result in more dramatic changes in fiber degrader abundance and activity in the gut microbiome. The increase of fiber from a diverse set of dietary foods, rather than single fiber supplements, may also contribute to increased consumption of other micronutrients and vitamins that affect the microbiome as well (24).

In this study, we sought to answer three questions: 1) does a diet rich in fiber from whole foods alter the overall microbiome? 2) does the intervention alter the abundance and diversity of known fiber-degraders (e.g., *Bifidobacterium*)? and 3) if we observe compositional shifts in the microbiome, do these correspond with metabolic changes in the production of short-chain fatty acids? To address these questions, we developed and employed a course-based undergraduate research experience (CURE) at UC Irvine to assess individual responses to a high-fiber diet (25). Integrating authentic research experiences within lab courses in order to facilitate a deeper understanding of academic and industrial research continues to be a priority for both national education reform and the American Society for Microbiology (25–28). During the intervention, participants were given ten meals each week from a food service that specializes in providing high fiber, unprocessed meals. Individuals tracked dietary information of macronutrients for every meal for three weeks, with the goal of increasing dietary fiber intake to 50 grams/day during a two-week intervention period. We then compared overall bacterial composition using metagenomic sequencing and assessed the production of volatile SCFAs using mass spectrometry. In addition to the shotgun metagenomic sequencing, we targeted a known-fiber degrader, *Bifidobacterium*, by analyzing its diversity using amplicon sequencing of the *groEL* marker gene, enabling a unique high-resolution view of the impact of a dietary fiber intervention on a key taxon.

## METHODS

### Study Design

Twenty-six UC Irvine students and instructors volunteered for a three-week high fiber diet intervention study (**Figure 1A**); only 22 individuals elected to provide stool samples for microbiome analyses (20 of whom we recovered enough sequence data for analysis, see **Supplemental Table 1**). The dietary intervention was approved by UC Irvine IRB # 2018-4297. For the first week of the study, all participants consumed their normal diets, tracking all nutritional information using the smartphone application MyFitnessPal (MyFitnessPal, Inc.). Prior to the end of week one, each subject provided three fecal samples from three days within the first week. The intervention commenced in week two, when participants were instructed to raise their dietary fiber intake to approximately 40 grams per day. To assist with the dietary shifts, we provided 10 meals per week with ∼15 grams of fiber ∼5.8 unique fruits or vegetables per meal from the food delivery service Thistle (https://www.thistle.co/ San Francisco, California, USA). During week three, subjects were encouraged to further increase fiber intake to ∼50 grams of fiber per day. Subjects provided three fecal samples from three days during week three, concluding the intervention period. As part of the CURE course, students were educated on human health, dietary information on high-fiber meals, the human gut microbiome, and the quantitative methods for microbiome analyses (from DNA extraction and library preparation to metagenome and statistical analyses), as previously described (25).

**Figure 1:**
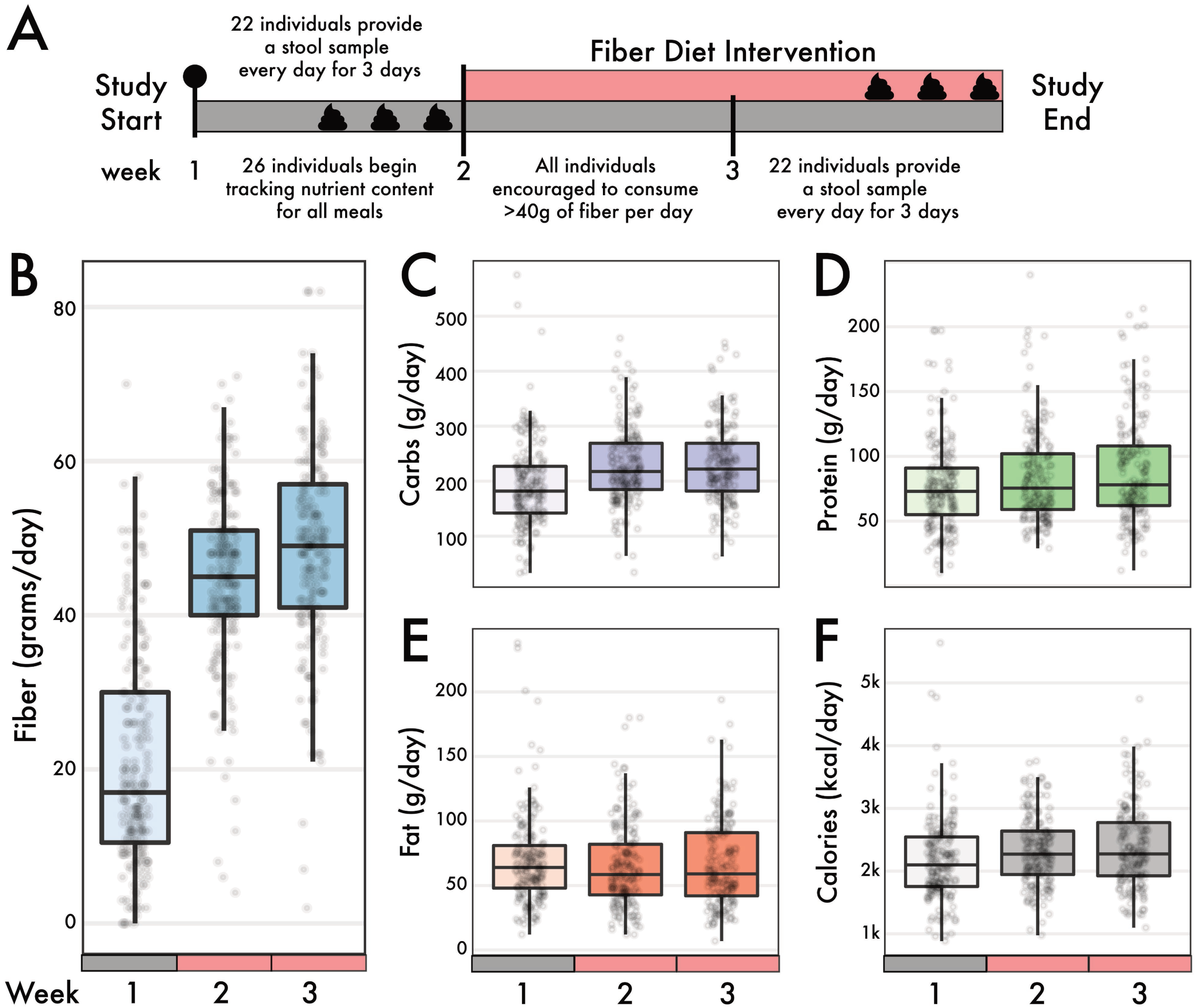
Intervention timeline and sample collection. The study subjects began eating their normal diet for one week, tracking all their food intake using the MyFitnessPal app. At the end of the week, each individual provided a daily fecal sample on three different days. At the start of week two, subjects started a wholesome high fiber diet, getting at least 40 grams of fiber per day. During week three, subjects were encouraged to get 50 or more grams of fiber per day. At the end of week three, each subject provided a fecal sample on three different days. B-F) Self-reported macronutrients from individuals using the MyFitnessPal app. Change in macronutrients across the 3-week diet intervention for B) fiber, C) carbohydrates, D) protein, E) fat, and F) calories. Fiber changed the most in magnitude between pre-intervention intake and during the diet intervention (linear mixed-effects model, p < 0.001). There were modest, but significant, changes in carbohydrate, protein, and caloric intake, but not fat intake, across the same time interval.

### Sample collection

Subjects were given materials to collect fecal samples at home. Each stool sample was split into three 2ml tubes by the individual and immediately stored in the freezer. When convenient, students transported their anonymized and coded samples using cold packs and insulated boxes to a common lab freezer. Upon the conclusion of the intervention period (week 1 or 3), all samples were transported to a −20 °C freezer.

### DNA extraction and metagenomic library preparation

To characterize the bacterial community composition of the samples, DNA was extracted with the ZymoBIOMICS 96 DNA Kit (Product D4309) from Zymo Research using the manufacturer’s suggested protocol. Sequencing libraries were prepared using the Illumina Nextera kit and methods described in Baym et al. (29). Briefly, DNA was diluted to 0.5ng/µl and added to 0.25µl of Nextera enzyme and 1.25 µl of Tagmentation Buffer. This mixture was incubated at 55 °C for 10 minutes and then placed on ice for the remainder of the protocol. Barcodes were added using the Phusion polymerase (New England Biolabs) and excess adaptors were cleaned using AMPure XP (Beckman Coulter Life Sciences) magnetic beads. Quality and concentration were assessed using a Picogreen assay (ThermoFisher) and the distribution of fragment sizes was determined using a Bioanalyzer. These libraries were loaded onto the Illumina Next-Seq 500 at 1.8 picomolar concentrations and sequenced using Illumina’s mid-output kit for 75 bp paired-end sequencing, resulting in a total of 144,023,583 reads and an average of 1,425,976 reads / sample (max: 5,902,966; min: 7) (**Supplemental Table 1**).

### Amplicon library preparation

To characterize the genetic diversity of *Bifidobacterium* at a finer-genetic scale than could be assayed by metagenomics, we used genus-specific primers to target this group for sequencing (30). Sequencing libraries were prepared by setting up an initial 25 µl PCR reactions with AccuStart II PCR ToughMix (2x), the *groEL* forward primer (5’-TCGTCGGCAGCGTCAGATGTGTATAAGAGACAGTCCGATTACGAYCGYGAGAAGCT-3’, 20 µM), and the *groEL* reverse primer (5’-GTCTCGTGGGCTCGGAGATGTGTATAAGAGACAGCSGCYTCGGTSGTCAGGAACAG-3’, 20 µM). The initial PCR ran for 28 cycles 95°C for 30 sec, 60 °C 30 sec, 72 °C 50 sec followed by the addition of 0.5 µl of dual Nextera XT index (Illumina) to each sample proceeding with an additional 8 cycles 95 °C 30 sec, 60 °C 30 sec and 72 °C 50 sec. Amplicons were pooled based on visual quantification of the bands on an agarose gel and purified using magnetic Speed Beads The pool was run on a MiSeq PE 300 at University of California Irvine’s Genetic High Throughput Facility resulting in a total of 20,052,935 reads and an average of 185,675 reads/sample (max: 6,815,601; min: 155).

### SCFA extraction and measurements

SCFA extractions were done following the methods by Zhao et al. (2005) (31). One-hundred mg of fecal material was added to 1ml of HPLC grade water and vortexed for two minutes. Ten microliters of 6N HCl was added to the fecal slurry and vortexed briefly. This mixture was incubated at room temperature for 10 minutes with occasional shaking. Afterwards the mixture was centrifuged at 14,000g for 1 minute, and 400 µl of the supernatant was transferred to a new tube, which was then filtered through a 0.22 µm filter. An aliquot (200 µl) of this suspension was then transferred to a glass vial with a 0.2 ml vial insert and stored at −20 °C. When running the sample, 10 µl of an internal standard of 10mM ethyl butyrate was added to the extraction prior to the run. Before running each sample, the instrument was calibrated using a standard comprising 100 mg/l of acetate, propionate, isobutyrate, butyrate, isovalerate, valerate, and ethyl butyrate. Six samples were run on an Agilent 7890A gas chromatograph with dual column FID detectors. Two microliters per extracted sample were hand-injected on a stainless-steel column (2 meters × 3.2 mm) containing 10% SP-1000 and 1% H3PO4 on 100/120 Chromosorb W AW (Supelco, Inc., Bellefonte, PA, USA). The flow rate of the N2 carrier gas was 26.14 ml/min. Between sets of six samples the instrument was washed using water and phosphoric acid. Peaks were auto integrated using ChemStation v1.0 on a PC running Windows 2000 (Microsoft). A subset of samples (n = 44 from 8 individuals) were run in duplicate to examine technical variation (see coefficient of variation (CV) in **Table 1**), and the average CV was 55%.

**Table 1:** Abundances of four SCFAs in samples pre- and post-intervention

|  | PRE<br>(MEAN) | PRE<br>(SD) | PRE CV | POST<br>(MEAN) | POST (SD) | POST<br>CV |
| --- | --- | --- | --- | --- | --- | --- |
| **ACETATE** | 480 mg/L | 208 mg/L | 43 % | 525 mg/L | 244 mg/L | 47 % |
| **PROPIONATE** | 277 mg/L | 112 mg/L | 41 % | 288 mg/L | 175 mg/L | 61 % |
| **BUTYRATE** | 244 mg/L | 107 mg/L | 44 % | 257 mg/L | 176 mg/L | 68 % |
| **VALERATE** | 49 mg/L | 31 mg/L | 65 % | 43 mg/L | 31 mg/L | 72 % |

### Metagenomic sequence analysis

Raw shotgun metagenome sequences were filtered using Prinseq v0.20.4 (32) to remove sequences that had a mean quality score of 30 or less. Reads from human DNA were also removed by aligning the filtered reads to the human genome (hg38), using Bowtie2 v2.2.7 (33), and keeping the reads that failed to align. A total of 130,755,383 paired-end reads (average 1,294,607 non-human reads/sample) were retained and passed through MIDAS, which assigns taxonomy to short read data using a marker gene approach (34). Species counts per sample represent the average of 100 subsamples, rarefied to 900 sequences per sample using the EcolUtils (v0.1) package in R. Taxonomy was also assessed using IGGsearch (35). To analyze functional differences related to SCFA metabolism between high and low fiber treatment groups, HUMAnN3 (36) was used with default parameters. All pathways within the MetaCyc pathway class “Fermentation to Short-Chain Fatty Acids” were searched for within the HUMANnN pathway output, which resulted in nine pathways used for analysis (37). For genes related to carbohydrate breakdown, we translated reads using Prodigal (38) to predict open reading frames (ORFs) and searched all ORFs against the Pfam database (39) with hmmer/3.1b2 (40). Resulting PFAM annotations were then screened against the CAZyDB.07202017 (41) with Blast/2.8.1 (42) using alignments >70% amino acid identity and 30% coverage. Alpha diversity and PERMANOVA analyses were performed using the Vegan v2.5-6 (43) package in R (44). Non-metric multidimensional analysis was done using the metaMDS function in Vegan on Bray-Curtis distances. StrainPhlAn (45), under default parameters, was used to analyze strain-level variation within the metagenomes. To root the phylogenetic tree, *Prosthecochloris aestuarii* (accession: GCA_000020625) was used, and two reference genomes of *Eubacterium rectale* (accession: GCA_000209935 and GCA_001404855).

### GroEL amplicon analysis

We downloaded 780 genomes from the genus *Bifidobacterium* on the PATRIC database (46). All genomes were screened for completeness by searching for 21 single-copy ribosomal marker genes using Prodigal (38) and HMMer v3.1b2 (40) with an E value of 1 × 10^−10^. The remaining 578 genomes were used to create a multi-locus, concatenated phylogeny of the ribosomal marker genes with ClustalO v1.2.0 (47) to produce a 4272 amino acid alignment for phylogenetic analysis using RAxML v8.0.0 (48) under the PROTGAMMABLOSUM62 model for 100 replicates. Next, we parsed the filtered genomes for the *groEL* gene sequences by using 260 non-redundant gene sequences to build a *groEL* phylogeny under identical parameters to the whole-genome analysis. The *groEL* amino acid sequences, alignment, and phylogeny were used to construct BLASTp, HMMer, and pplacer reference databases for metagenomic analyses. For each *groEL* amplicon library, sequences were quality trimmed and adapters were removed with BBDuk (49) (qtrim=rl trimq=10 ktrim=r k=25). Paired end sequences were merged together with BBMerge (49) and, if paired reads did not overlap, only the forward read was retained. The reads were then searched against the *groEL* reference databases using BLAT (50) and hmmsearch, respectively. Passed reads were aligned with ClustalO to the pplacer reference package and placed onto the *groEL* reference phylogeny using pplacer v1.1.alpha17 (51). Relative abundance was calculated from the single branch assignments and aggregated at the species level to be normalized by the total number of extracted *groEL* gene sequences. We show that the phylogenetic relationship between species of *Bifidobacterium* based on the *groEL* gene closely reflects a phylogeny based on 21 single copy marker genes from 578 *Bifidobacterium* genomes (**Supplemental Figure 4)**.

### Statistical analysis

Permutational analysis of variance (PERMANOVA) was conducted on Bray-Curtis dissimilarities at the genus level with 999 permutations using the Adonis test in the Vegan package in R (see Data availability and GitHub). We tested the effect of the intervention (pre- versus post-fiber increase), the effect of the individual, and the interaction between these two factors. Genus contributions to significant results from the PERMANOVA model were determined by passing the resulting PERMANOVA object through the coefficients function found in the base Stats package R. A similar procedure was used to analyze compositional differences between CAZy enzymes and HUMAnN gene predictions in the metagenomes, with permutations on Euclidean distances. Linear mixed effects models, using the nLME package (44) in R, were also conducted for comparison because they take repeated measures into account. Specifically, to support the PERMANOVA analysis of beta diversity, an LME was performed on the rank-transformed first principal coordinate of a principal coordinates analysis on the Bray Curtis community dissimilarity matrix. Individual was used as the random effect and the model used the default autoregressive (Lag 1) structure (AR1) for regression across a time-series. For the functional analyses, reads analyzed using HUMAnN3 were normalized by copies per million; CAZy were normalized to the total number of reads per metagenome and compared using Wilcoxon rank sum test. Gene features for HUMAnN were reduced by analyzing only unstratified data, for which 70% of samples had non-zero reads mapping to each feature. HUMAnN pathway abundances were analyzed in their entirety with stratification and without feature reduction. Lefse (52) was used to determine pathways which may differentiate pre- vs post-intervention samples. Wilcoxon rank sum tests were also used to compare nutritional and gene differences between intervention periods when residuals were not normally distributed and reads or macronutrients were averaged out within individuals (by treatment) to account for repeated measures. When normality assumptions of residuals were met (tested using the Shapiro-Wilk test) ANOVAs were used. To assess which taxa were correlated with changing amounts of fiber, all species within each sample (the rarefied species abundance matrix) and fiber were correlated using the Corrr package v0.4.2 (53) in R. To analyze which genera co-correlate with the genus *Bifidobacterium*, Spearman correlations were used and, where appropriate, p-values were corrected (q-value) for multiple comparisons using a false discovery rate cutoff of 0.05. To assess significance of strains between individuals, cophenetic distances were calculated on the RAxML tree output from StrainPhlAn and passed into the above PERMANOVA model.

### Data availability

All scripts are stored on GitHub (https://github.com/aoliver44/Fiber-Analysis). All metagenomic and amplicon sequences are available on NCBI under the Bioproject PRJNA647720. Metadata linking the shotgun metagenomes and *groEL* sequences with the appropriate sample ID and intervention can be found in Supplementary Table 1.

## RESULTS

### Dietary intervention within the CURE course

Twenty-six individuals participated in a CURE course at UC Irvine, designed to tandemly investigate pedagogical methods (25) and the role of fiber on the microbiome. We collected nutritional data from all 26 individuals who initially began the intervention, over three weeks (one week prior to, and two during, the dietary intervention) (**Figure 1A**). We collated the total amount of macronutrients consumed per day, including fiber, protein, carbohydrates, fats, as well as overall calories (**Figure 1B-F**). Additionally, we informally surveyed food items the study participants frequently used to supplement their meal plans, beyond the meals supplied from Thistle, and found that items such as fiber fortified cereals, lentils or beans, and berries were common (25). For the intervention, subjects increased their average fiber consumption from 21.0 g/day (± 14.2 g/day) before the intervention to 46.4 g/day (± 12.5 g/day) during the intervention (**Figure 1B**; Wilcoxon rank sum test, p < 0.0001). While these dietary shifts increased carbohydrate intake by an average of 84% (36 g) during the intervention (p = 0.013), other macronutrients measured, such as calories, fat, and proteins, did not significantly change (p > 0.05) (**Figure 1C-F**).

### Diet intervention altered gut microbial community composition within individuals

To evaluate whether increased fiber consumption contributed to shifts in the gut microbiome, we characterized the microbial communities from 20 individuals using 86 shotgun metagenomic libraries collected before and after the fiber intervention (**Figure 2A**). Alpha-diversity of microbial taxa decreased during the high fiber diet intervention as measured by the Shannon diversity index (**Figure 2B**)(Wilcoxon rank sum test, p < 0.05). Using alternative approaches to assess taxonomy and diversity (see methods) showed either no change or supported the decreasing trend of diversity during the intervention period (**Supplemental Figure 1).**

**Figure 2:**
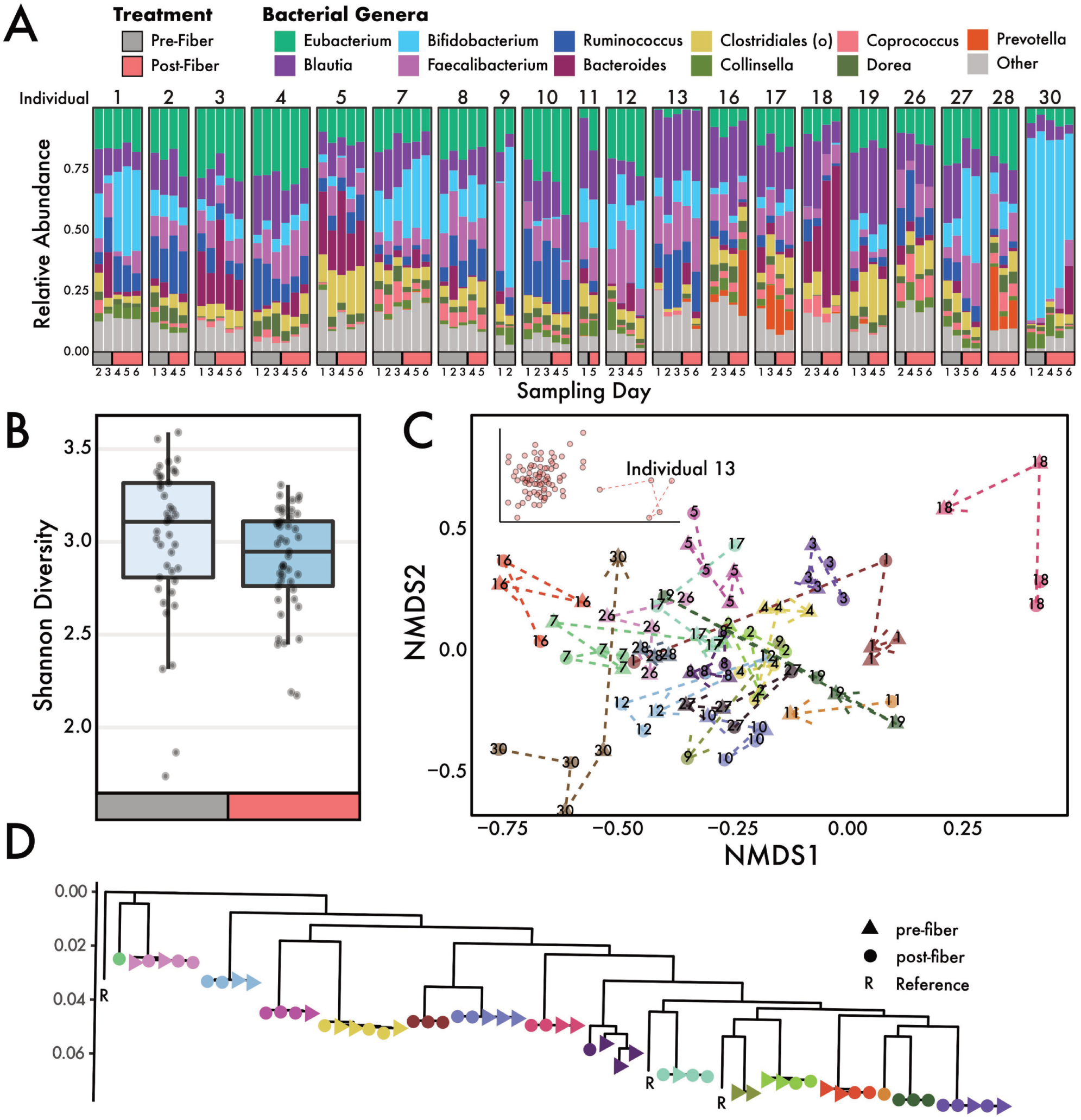
Microbiome community composition through a dietary fiber intervention. A) Relative abundances of genera detected in microbiomes from individuals throughout the diet intervention study. B) Alpha diversity, measured using the Shannon index, changed significantly during the intervention period (Wilcoxon, p < 0.05). C) NMDS ordination showed that samples from individuals mostly group together. Dotted lines connect the same individual and point towards the final post-fiber intervention sample. Samples in this study were highly personalized: the individual explained 78% of the variation in the data. The inset shows an extended version of the NMDS plot that includes Individual 13. D) A phylogeny of *Eubacterium rectale* strains found in individuals (denoted by color) during the intervention.

Despite little difference in alpha-diversity, beta-diversity changed significantly in response to a high fiber diet. Multivariate analysis of marker gene abundances showed that most of the variation in microbiome composition could be explained by the individual (PERMANOVA: main individual effect: R^2^ = 0.78, p < 0.001, **Supplemental Table 2**). The diet intervention shifted the microbial composition of the entire study cohort significantly (main intervention effect: R^2^ = 0.014, p < 0.001). Within samples from each individual, the pre- and post-diet intervention samples explain significant variation in the community composition (intervention-by-individual effect: R^2^ = 0.083, p < 0.001). A linear mixed-effects (LME) model confirmed these results, which identified diet as a significant determinant of an individual’s microbiome composition (LME, p < 0.01). Individual gut microbiome samples grouped together in nonmetric multidimensional space (nMDS; **Figure 2C),** further providing support that each individual is associated with a unique microbiome. Some individuals (i.e., Individual 13) gut microbiomes were more distinct from others (**Figure 2C inset)**. Additionally, we used *Eubacterium rectale* (due to its high coverage in our data) to ask whether the diet intervention had an impact at the strain level. Strains were highly individual specific (PERMANOVA: main individual effect: R^2^ = 0.99, p < 0.001) and did not change in response to increased fiber intake (p > 0.05; Figure 2D).

We next parsed the taxonomic data to assess which microbial taxa increased or decreased in response to the diet intervention. One species in the family *Lachnospiraceae* was significantly negatively associated with increasing fiber intake (Spearman, r = −0.43, q = 0.01) (**Supplemental Figure 2A, B**). *Coprococcus sp.* and *Anaerostipes hadrus* were both positively associated with increasing fiber intake, but this association was not significant when p-values were FDR-corrected for multiple comparisons (r = 0.32, q = 0.33 both species) (**Supplemental Figure 2A**). Furthermore, positive linear coefficients of a PERMANOVA model, which detect differences between community composition due to the diet intervention, included genera such as *Bifidobacterium, Bacteroides*, and *Prevotella* (**Figure 3A**). Conversely, *Blautia* and *Ruminococcus* contributed negative linear coefficients to the PERMANOVA model (**Figure 3A**).

**Figure 3:**
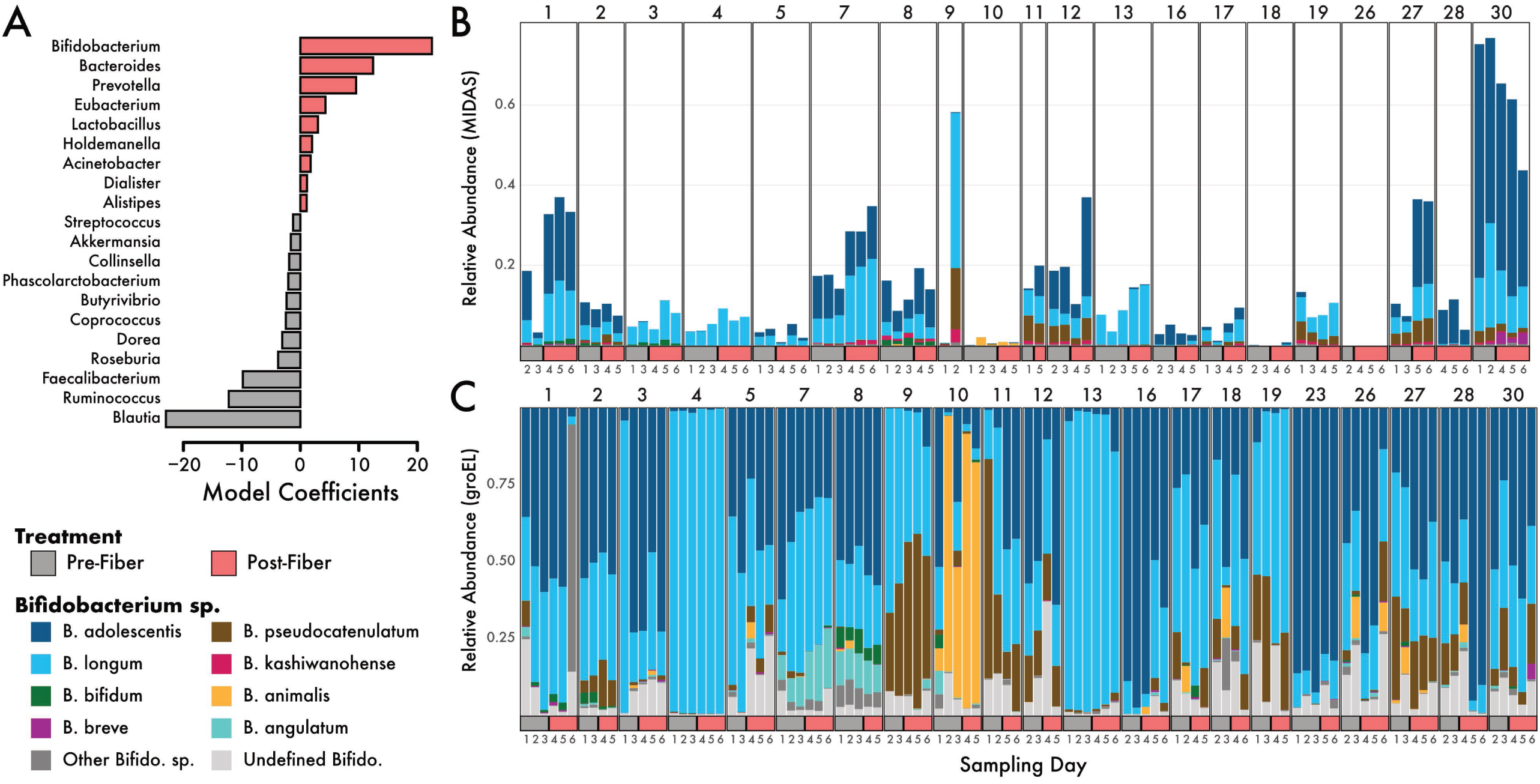
GroEL amplicon analysis of *Bifidobacterium* during the fiber intervention. A) Model coefficients of the PERMANOVA analysis (model: species ∼ Individual^*^Intervention). Species with high coefficients (positive or negative) were best able to distinguish the pre vs post diet intervention groups. Only the top 20 genera are shown. The genus Bifidobacterium had the largest positive coefficient, indicating that it was important to the model for disguising microbiomes before and after the diet intervention. Relative abundances of 12 detected species of Bifidobacterium from B) shotgun metagenomics and C) groEL amplicon sequencing.

### *Bifidobacterium* species were enriched by the diet intervention

Of the 105 microbial genera detected in this study, *Bifidobacterium* was the strongest predictor genus for the post-intervention microbiomes (**Figure 3A).** Indeed, taxonomic analysis of the metagenomic samples identified *Bifidobacterium* abundances increasing, on average, 1.4-fold between the pre- and post-intervention periods (**Supplemental Figure 3A)**. Further, we identified several species of *Bifidobacterium* present within and across individuals, with *B. adolescentis* being the most abundant species on average (**Figure 3B).** When we investigated the taxonomic profiles at the species level, we found that *B. adolescentis, B. biavatii, B. breve, B. longum*, and *B. ruminantium* all increased in mean abundance on a high fiber diet whereas the other, lesser abundant species exhibited no change or decreased in abundance (**Supplemental Figure 3B**).

Given that *Bifidobacterium* was the strongest predictor genus in the post-fiber gut microbiomes, we employed a targeted analysis into the diversity within *Bifidobacterium* to examine species-level patterns. Specifically, we applied targeted amplicon approaches to amplify the *groEL* gene, a conserved phylogenetic marker gene to track *Bifidobacterium* diversity (**Supplemental Figure 4)**. Using phylogenetic inference of the *groEL* gene, we compared the observed *Bifidobacterium* diversity observed at the community level to our targeted analysis of the *groEL* gene. Similar to the metagenomic analysis, we found that individuals were largely comprised of *B. adolescentis* and *B. longum*, with six other abundant species of *Bifidobacterium* (**Figure 3C).** This analysis also revealed extensive *Bifidobacterium* diversity within the human gut, detecting 22 species across all individuals.

Since *Bifidobacterium* species are known to participate in cross-feeding with other gut microbes (reviewed in (54)), we next assessed the co-occurrence of *Bifidobacterium* with other genera. *Bifidobacterium* was positively correlated (r = 0.43, q = 0.001) with an increasing abundance of *Lactobacillus* and negatively correlated with *Roseburia* (r = −0.49, q = 0.0002) and *Ruminococcus* (r = −0.38, q = 0.007) (**Supplemental Figure 5**) suggesting possible species interactions between these taxa.

### Genes involved in inositol degradation increase on high fiber diet

Our results demonstrate that a shift in dietary fiber consumption influenced compositional changes in the gut microbial community. As such, we sought to correlate the observed taxonomic shifts to functional shifts, particularly the enrichment of genes related to carbohydrate degradation. Despite taxonomic shifts at the individual level, we observed no changes in in the overall abundance (average number of normalized reads) mapping to gene families for glycoside hydrolases (GH) (Wilcoxon, p = 0.42), carbohydrate esterases (p = 0.58), glycoside transferases (p = 0.73), and polysaccharide lyases (p = 0.77) as a result of the intervention (**Supplemental Figure 6A**). No individual families GH and polysaccharide lyase CAZy classes changed in abundance during the intervention when corrected for multiple comparisons (Wilcoxon, p > 0.05, **Supplemental Figure 6B**). Further, the diversity (**Figure 4A,** n = 106 families, ANOVA, p > 0.05) and composition (PERMANOVA, p > 0.05) of GHs detected were indistinguishable between the pre- and post-intervention samples. Compositional analysis all genes identified by HUMAnN revealed no individual signature (PERMANOVA: main individual effect: R^2^ = 0.017, p > 0.05, **Figure 4B**), and no shifts in response to a high fiber diet (intervention-by-individual effect: R^2^ = 0.015, p > 0.05). We performed a linear discriminant analysis to determine if there were pathways that were differentially abundant due to the diet intervention, and found inositol degradation (in addition to several unintegrated pathways) to be increased in abundance on a high fiber diet (**Figure 4C, Supplemental Figure 7**). For the pathways involved in SCFA metabolism, we found no significant (Wilcoxon, p > 0.05) changes as a result of the high fiber diet (**Figure 4D**).

**Figure 4:**
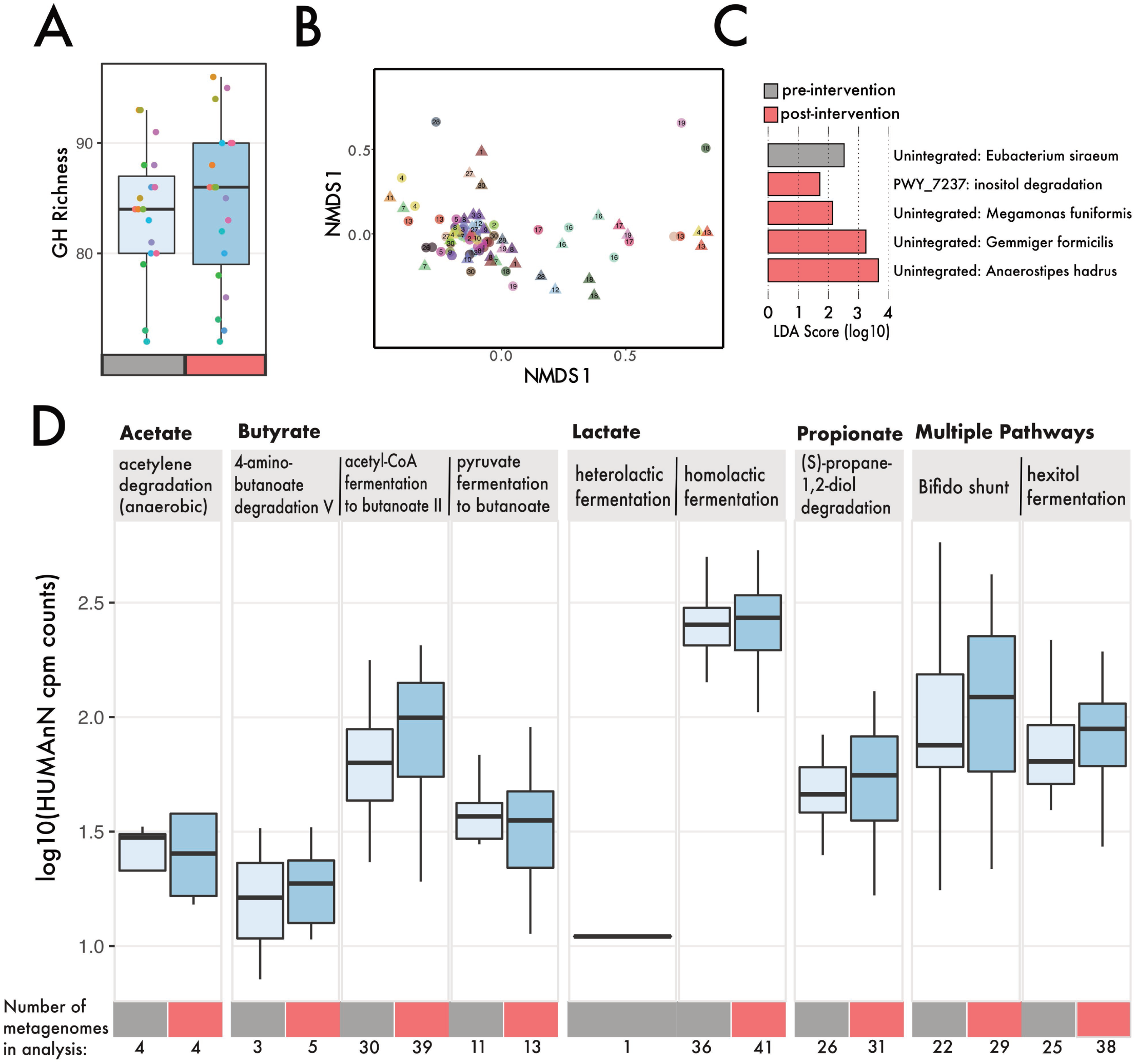
Genes involved in carbohydrate degradation and SCFA metabolism within metagenomes. A) Number of distinct glycoside hydrolase families within individual metagenomes (different colored circles), separated by pre-intervention (mean = 83) (grey) and post intervention (mean = 84) (red). B) NMDS ordination of Euclidean distance matrix based on 19680 gene features, shape denotes intervention (triangle = pre-, circles = post-) and individuals are separated based on color. C) Lefse analysis of pathways that differentiate samples by intervention. D) Log abundance (copies per million) of pathways involved in SCFA production.

### Fecal short-chain fatty acid concentrations were unaltered by the diet intervention

While the presence of genes related to SCFA production provide insights into the functional changes of the microbiome, these genes only reflect the genomic potential to process these pathways. Therefore, we applied a targeted GC/MS analysis on 149 samples from 18 individuals for the presence of SCFA molecules. Across the intervention period, the average abundance of acetate, propionate, butyrate, and valerate increased (Table 1); however, these increases were not statistically significant (LME, p > 0.05) (**Figure 5)**. For the eight individuals with samples run in duplicate, three had biological differences that were greater than the technical variation seen in the duplicates (**Supplemental Figure 8**). Acetate had the least technical variation (mean CV = 45%), followed by propionate (mean CV = 51%) (**Table 1, Supplemental Figure 8**).

**Figure 5:**
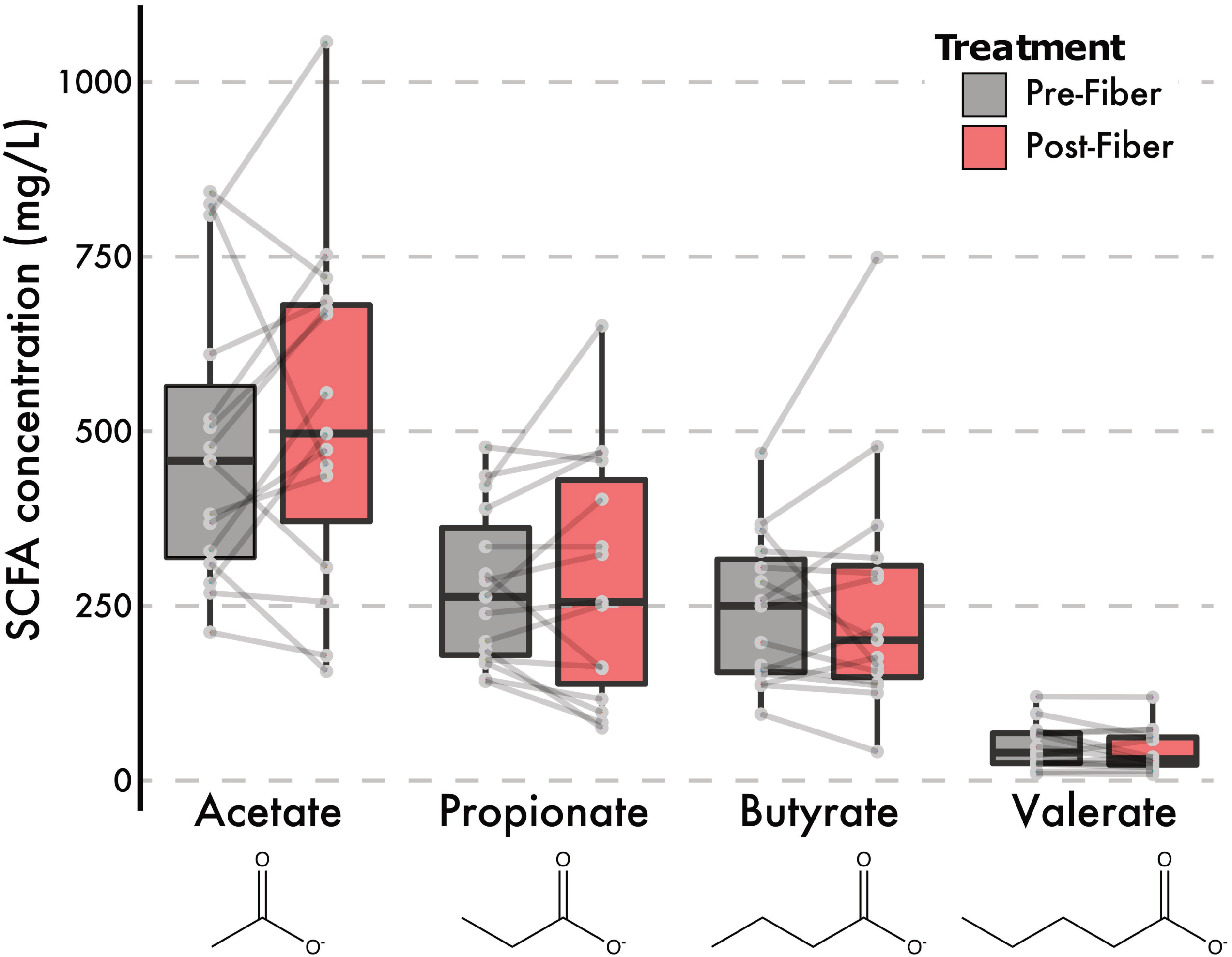
GC-FID measurements of fecal volatile SCFAs during intervention. Fecal SCFA abundances, averaged across replicates where applicable, before and after the intervention.

## DISCUSSION

We examined the impact of dietary foods, rich in their diversity of fiber, on the human gut microbiome. We expected that an increase in fiber consumption through whole foods consumed would lead to a more generalizable shift of the microbiome in contrast to previous studies that utilize a single fiber supplement. For instance, a recent meta-analysis (55) found mixed results in how fiber may impact the gut microbiome richness and composition. Among papers published prior to October 2017, only 18% (12 out of 64) studies (56, 57, 66, 67, 58–65) contained food-based fiber interventions, and most of these studies only modified one aspect of diet (e.g., addition of whole grain breakfast cereal). One study in particular increased dietary fiber by 40g from a diverse set of foods during a five day period (57). The authors similarly found microbiome composition changes within individuals when they accounted for differences in the subjects’ starting microbiomes. Despite the variation in implementing a fiber intervention, it is becoming increasingly clear that fiber alters the composition of the gut microbiome (17) and the associated microbial changes affect human health (i.e., type 2 diabetes mellitus (3)). A common observation in fiber intervention studies (55) is the specific involvement of the genus *Bifidobacterium* in response to fiber interventions. However, to our knowledge, no study has documented how fiber impacts the genus at the strain-level in the human gut.

### Does a diet intervention rich in fiber alter the microbiome?

Past studies have shown that an increase in the diversity of dietary foods could lead to an increase in microbial diversity (24). Moreover, individuals living in rural societies often harbor far greater gut microbial diversity than individuals from western societies (68–70), which may in part be linked to a greater proportion of plant-based polysaccharide intake. However, we did not measure an increase in species diversity (alpha diversity) after subjects consumed >40g of fiber from a diverse set of foods (**Figure 2B**). These results could be attributed to the brevity of the intervention as the rapid change in dietary composition may result in the loss of microbes poorly adapted to recalcitrant carbohydrates. Similarly, other studies have reported finding no increases in alpha diversity as a result of a fiber intake (57, 71–74), which may indicate a trade-off where fiber-degraders increased while other taxa decreased. Although alpha diversity was unaffected, we did observe a significant impact of the high-fiber diet on microbial community composition (beta diversity) (**Figure 2**). The composition of microbial communities within individuals shifted ∼8% during the intervention period. We found changes in communities to be at broader taxonomic levels than strain-level. We were able to examine strains of *E. rectale* due to its high coverage in our data, and showed these strains stayed constant and individual specific during the intervention (**Figure 2D**). Future work should determine if this pattern holds up for other species. While we suspect the high fiber diet treatment played an instrumental role in shifting the microbial composition, we cannot rule out other factors such as host genetics or non-dietary behaviors. As discussed, many food-based fiber interventions have shown mixed results on changing the microbial communities (6, 75). The drastic increase in fiber from a variety of foods may lead to rapid shifts in community composition over the two-week period. Changes in community composition pre- and post-intervention were largely driven by shifts in known-fiber degraders, such as *Bifidobacterium, Bacteroides*, and *Prevotella* (**Figure 3A**).

We expected the taxonomic shifts in the microbiota would be associated with changes in the functional potential of the microbial communities (**Figure 4**). While we initially hypothesized that a high fiber diet would increase the abundance or diversity of carbohydrate active enzymes, we did not detect changes associated with the intervention (**Supplemental figure 6)**. Our findings support a similar result showing no difference in CAZy abundance due to increased fiber intake (74). We acknowledge that sequencing depth is an important consideration in the detection of genes; increasing reads beyond our ∼1.3 million paired-end reads (avg per sample, **Supplemental table 1**) may allow for greater detection. However, we did find a notable increase in the abundance of genes mapping to the inositol pathway (**Figure 4C**). We suspect that the increased consumption of fiber-fortified cereals and legumes, which contain higher levels of inositol, during the diet intervention allowed for an expansion in organisms capable of breaking down this sugar. There is substantial interest in the role of inositol (specifically phytic acid) in its protective role against colon cancer and other metabolic disorders (76, 77). Next, we assessed whether genes involved in SCFA metabolism changed in abundance during the intervention. Although appreciable cross-feeding between lactate-producing *Bifidobacterium* spp. and butyrogenic bacteria has been shown (78) we did not find significant increases in genes involved in various SCFA metabolic pathways (**Figure 4D**). This further supports our results showing no clear correlations between *Bifidobacterium* spp. and butyrate-producers within our diet intervention (**Supplemental figure 5)**. Indeed, we would not be the first to suggest that perhaps these complex trophic interactions require more time to establish (17). Rather, our results suggest that while broad taxonomic shifts occur, these do not correspond to changes in functional potential and fine-scale (intraspecies) shifts are less susceptible to dietary shifts on short-term timescales.

### Does the intervention alter the abundance and diversity of Bifidobacterium, a known fiber-degrader?

Many studies have indicated that bifidobacteria (often identified as the genus *Bifidobacterium* by FISH probes, PCR, or DNA-sequencing) are highly abundant in the gut following increased fiber intake (meta-analysis of 51 studies (55)). Increased abundance of *Bifidobacterium* is somewhat unsurprising, as they harbor numerous genetic components, such as carbohydrate active enzymes, that make them especially adapted to a fiber-rich diet (79). In one study, both resistant potato starch and inulin increased the relative abundance of *Bifidobacterium* spp.; however, the 16S amplicon sequencing in this study did not have the resolving power to identify which species of *Bifidobacterium* were increasing (17). Using a targeted amplicon approach, the *groEL* gene, has been shown to delineate species of *Bifidobacterium* that otherwise share >99% sequence identity in the 16S rRNA gene, making it a robust marker gene for analyzing within-genus species diversity (80). In our study, the most abundant species of *Bifidobacterium* were *B. adolescentis* and *B. longum*, both of which are efficient degraders of plant-based fructo-oligosaccharides (FOS) and produce acetate and lactate in the process (81). Mirroring our results, other studies have found selective increases in certain species of *Bifidobacterium* as a result of carbohydrate intake; for example, in one study, intake of inulin resulted in a greater increase of *B. adolescentis* (82). We speculate that on a high fiber diet, bifidobacteria are the initial members of the community accessing fiber substrates, easily adapted to utilize various FOS, and pivotal to the creation of the initial metabolic cross-feeding networks. Future studies should extend the intervention period to examine the dynamics of longer-term trophic interactions in response to increased dietary fiber intake.

### Can we detect diet-induced changes in the abundance of fecal short-chain fatty acids?

While SCFAs did generally increase during the diet intervention, trending toward their naturally occurring gut ratio of 3:1:1 (acetate:propionate:butyrate) (12, 83, 84), we did not observe a statistically significant increase in SCFAs post-intervention. Static fecal concentrations of SCFAs may not reflect the total pool of molecules fluxing through a given individual, as the molecules are preferred substrates of the cells lining the gut epithelia (15). It is also possible that the intervention period was too short to observe increases in SCFA abundances.

It should be noted that accurate SCFA measurements are notoriously difficult. Our examination of technical variability within 44 samples from eight individuals showed that technical variation between pre-intervention replicates or post-intervention replicates was greater than the average difference between pre- and post-intervention for any given SCFA. One study reported high intra-fecal variability of butyrate quantification (coefficient of variation = 38%), prior to optimizing a freeze-drying method (85). Numerous studies have indicated the benefit of SCFAs to human health (5, 86); yet the heterogeneity in reported acetate, propionate, and butyrate abundances remains high. In one meta-analysis of fiber studies, only butyrate was generally found to increase with fiber intake, yet the heterogeneity of reported results was 70% (*I^2^*), similar to other SCFAs analyzed (55). Outside of technical limitations, shifts in microbial community structure are not predictive of changes in static measurements of fecal SCFA abundances (87). The difficulty of finding meaningful correlations between microbiome composition and SCFA abundances likely reflects a failure to measure both circulating and fecal SCFAs across time in conjunction with microbial abundances. Indeed, it has been observed that fecal levels of acetate are inversely related to the rate of its absorption (88). Future studies are needed to confirm whether correlation analysis between fecal SCFAs and microbiome composition is a useful tool to understand the interplay between microbiome, SCFAs, and health.

In sum, our results indicate that gut microbial communities are malleable to an influx in recalcitrant carbohydrates, contributing to significant community and functional shifts in certain metabolic pathways. However, these compositional changes did not correspond to broad functional changes, at least over the short-term timescales for this intervention. Further studies exploring the impact of timing and composition of dietary fiber interventions, particularly while taking into account the starting composition of the gut microbiomes of study participants, are critical for understanding the generalizability of fiber interventions for engineering microbiomes. Increasing fiber intake could have the most impact in contexts where low gut microbial diversity increases risk of *C. difficile* infection, such as for nursing home residents, cancer patients or after antibiotic treatment.

## ACKNOWLEDGMENTS

We would like to acknowledge the T32 training grant which supported Andrew Oliver (1T32AI14134601A1) from UC Irvine’s training program in microbiology and infectious diseases. We would also like to acknowledge the UCI Microbiome Initiative for supporting the study, Thistle, for supporting our provision of high fiber meals, and Heather Maughan for thoughtful edits to the manuscript. We would like to give an enormous thank you to the students from the course M130L at UC Irvine Spring in 2018.

## TABLE LEGENDS

**Supplemental Table 1:** Sequence statistics, accession IDs and experimental metadata.

**Supplemental Table 2:** PERMANOVA model and term results.

## FIGURE LEGENDS

**Supplemental Figure 1:** Comparisons of diversity measures obtained using different databases for taxonomic assignments. IGG_rich and MIDAS were run using default parameters. IGG-Lenient was run at 25% species quality and 15% marker genes.

**Supplemental Fig 2:** A) Correlations above a r > 0.2 cutoff of microbial abundance and fiber intake. Only *Lachnospiraceae* bacterium 51870 was significantly negatively correlated at an FDR cutoff of 0.05. B) Raw spearman correlation of a species of *Lachnospiraceae* with fiber intake.

**Supplemental Figure 3:** A) Mean abundance (MIDAS read counts) of the genus Bifidobacterium during the diet intervention. Points are colored by individual. B) Changes in mean abundance of each species of Bifidobacterium, detected by MIDAS, during the diet intervention period.

**Supplemental Figure 4:** *Bifidobacterium* phylogenetic analyses. A) Multi-locus phylogenetic analysis of conserved ribosomal marker genes. B) Phylogenetic analysis of the *groEL* gene sequences used for amplicon analyses. The top 8 species observed are shown.

**Supplemental Figure 5:** Significant (FDR = 0.05) correlations from comparing abundances of 99 different genera with *Bifidobacterium.*

**Supplementary Figure 6**: A) Average abundances (normalized reads) of carbohydrate active enzymes within individuals during the diet intervention period. Each different color point represents an individual, and lines connect the same individual. B) Log2 transformed GH and polysaccharide lyase gene abundances during the intervention.

**Supplemental Figure 7:** Inositol degradation abundance (normalized copies per million), for metagenomes before and after the intervention.

**Supplemental Figure 8:** Technical variation of SCFAs seen in a subset of samples run in duplicate. A) Amount of SCFAs by individual, with color denoting if it was measured during replicate 1 or 2. B) Normalized (by mean) difference between absolute difference between treatment, subtracted by absolute difference between technical replicates. Larger negative values suggest differences between technical replicates were larger than the differences detected between pre- and post-intervention arms.

